# Global connectivity and function of descending spinal input revealed by 3D microscopy and retrograde transduction

**DOI:** 10.1101/314237

**Authors:** Zimei Wang, Brian Maunze, Yufang Wang, Pantelis Tsoulfas, Murray G. Blackmore

## Abstract

The brain communicates with the spinal cord through numerous axon tracts that arise from discrete nuclei, transmit distinct functions, and often collateralize to facilitate the coordination of descending commands. In efforts to restore supraspinal connectivity after injury or disease, this complexity presents a major challenge to interpreting functional outcomes, while the wide distribution of supraspinal nuclei complicates the delivery of therapeutics. Here we harness retrograde viral vectors to overcome these challenges. We demonstrate highly efficient retrograde transduction by AAV2-Retro in adult female mice, and employ tissue clearing to visualize descending tracts and create a mesoscopic projectome for the spinal cord. Moreover, chemogenetic silencing of supraspinal neurons with retrograde vectors results in complete and reversible forelimb paralysis, illustrating effective modulation of supraspinal function. Retrograde efficacy persists even after spinal injury, highlighting therapeutic potential. These data provide a global view of supraspinal connectivity and illustrate the potential of retrograde vectors to parse the functional contributions of supraspinal inputs.

## SIGNIFICANCE STATEMENT

The complexity of descending inputs to the spinal cord presents a major challenge in efforts deliver therapeutics to widespread supraspinal systems, and to interpret their functional effects. Here we demonstrate highly effective gene delivery to diverse supraspinal nuclei using a retrograde viral approach and combine it with tissue clearing and 3D microscopy to map the descending projectome from brain to spinal cord. These data highlight newly developed retrograde viruses as therapeutic and research tools, while offering new insights into supraspinal connectivity.

## INTRODUCTION

Descending axon tracts in the spinal cord originate from diverse nuclei throughout forebrain, midbrain, hindbrain and serve a wide range of motor, sensory, and autonomic functions (Kim et al., 2017; Liang et al., 2011). Coordinated activity in multiple tracts often converges on the same motor output. For example, distinct phases of targeted forelimb movement are controlled by different sets of corticospinal tract distributed through motor cortex, in coordination with other nuclei including the red nucleus and reticular formation (Alstermark et al., 2004; Esposito et al., 2014; Kuypers H. and Martin, 1982; Lemon, 2008; Siegel et al., 2015; Wang et al., 2017). Clarifying the different contributions of supraspinal populations to basic motor control, remains a challenge. In addition, this complexity presents a significant challenge for the study and the treatment of spinal injuries. Spinal injuries in both human patients and animal models are highly variable, leaving each individual with different subsets of descending axon tracts intact. Thus, a major question in the field is how plasticity in distinct supraspinal tracts, both spontaneous and treatment-induced, yields behavioral improvements. Conversely, the fact that supraspinal neurons are distributed so widely and carry both discrete and overlapping functions presents a challenge for the delivery of therapeutics. Genetic interventions that boost the intrinsic growth ability of supraspinal neurons are showing promise in enhance regenerative axon growth from various types of neurons including corticospinal, rubrospinal, and reticulospinal projections (Blackmore et al., 2012; Geoffroy et al., 2016; Liu et al., 2010, 2017; Z. Wang et al., 2015; Wu et al., 2017). In these proof-of-principle studies vectors are typically delivered by direct brain injection, yet this approach is impractical to reach all supraspinal neurons in rodent models, let alone human patients.

The development of new viral vectors with effective retrograde transduction properties affords new opportunities to address these challenges. Descending axons converge to relatively small target fields in the spinal cord, and thus a small number of retrograde injections could potentially affect widely distributed supraspinal neurons of origin (Frampton et al., 2005; Nassi et al., 2015). The potential utility of this retrograde approach is well recognized, but has been limited by inadequate efficiency of transduction, particularly for corticospinal tract neurons (Jara et al., 2012; Klaw et al., 2013). Here we harness a recently developed adeno-associated serotype termed AAV2-Retro as a tool to clarify supraspinal connectivity and function (Tervo et al., 2016). We first demonstrate that AAV2-Retro delivered to the cervical and lumbar spinal cord drives rapid and highly efficient transduction of supraspinal populations including reticular, red nucleus, and corticospinal. Importantly, retrograde transduction remained equally effective when vectors were delivered after spinal injury, highlighting the potential for therapeutic delivery. We then employ tissue clearing and light-sheet microscopy to present detailed visualizations of supraspinal neurons and descending axon tracts in whole brain preparations. Finally, we test a retrograde viral approach to probe the function of supraspinal inputs. We found that after retrograde delivery of inhibitory Designer Receptors Exclusively Activated by Designer Drugs (DREADDs), receptor activation in transduced supraspinal populations was sufficient to induce complete forelimb paralysis. Moreover, when we employed transgenic targeting in conjunction with Retro-DREADD delivery to enrich expression in corticospinal tract neurons, receptor activation produced movement deficits highly consistent with selective CST ablation. Overall these data highlight the utility of a retrograde viral approach for therapeutic gene delivery, as a tool for neuroanatomical tracing, and for functional dissection of supraspinal circuits.

## MATERIALS AND METHODS

### Viral injection and spinal injuries

All animal procedures were approved by the Marquette University Institutional Animal Care and Use Committee and complied with the National Institutes of Health Guide for the Care and Use of Laboratory Animals. AAV2-Retro-CAG-tdTomato, AAV2-Retro-CAG-EGFP, and AAV2-Retro-CAG-CRE were purchased from the University of North Carolina Viral Vector Core. AAV2-Retro-hSyn-DIO-hM4D(Gi)-mCherry (44362) and AAV2-Retro-hSyn-hM4D(Gi)-mCherry (50475) were purchased from Addgene. Adult female C57/Bl6 mice (>8wks age, 20-22g) were anesthetized by Ketamine/Xylazine and the cervical spinal column exposed by incision of the skin and blunt dissection of muscles. Mice were mounted in a custom spine stabilizer. Viral particles were mixed with Cholera Toxin Subunit B conjugated to Alexa Fluor 488 (CTB-488) in sterile 0.9% NaCl (C22841-Thermofisher, Waltham, MA, final concentration of 2%), and were injected to the spinal cord through a pulled glass micropipette fitted to a 10μl Hamilton syringe driven by a Stoelting QSI pump (catalog # 53311) and guided by a micromanipulator (pumping rate:0.04ul/min). Viral/CTB mixtures were injected between C4 and C5 vertebrae, bilaterally, 0.35mm lateral to the midline, and to depths of 0.6mm and 0.8mm. For lumbar injections, the T9 vertebrae was used as a landmark to identify the L3 and L4 vertebrae. Injections were made between L3 and L4 vertebrae, bilaterally, 0.35mm lateral to the midline, 0.6mm and 0.8mm depth. Cervical dorsal hemisections were performed as in (Blackmore et al., 2012; Zimei Wang et al., 2015). Briefly, mice were anesthetized, their spinal cord exposed, and mounted in a custom spine stabilizer as described above. Using a Vibraknife device (Zhang et al., 2004), in which a rapidly vibrating blade is controlled via a micromanipulator, a transection was made between the 4^th^ and 5^th^ cervical vertebrae, extending from the midline beyond the right lateral edge of the spinal cord, to a depth of 0.85mm. Injections were performed immediately after injury, unilaterally to right spinal cord, 0.3mm rostral to the injury, 0.35mm lateral to the midline, and to 0.6mm and 0.8mm depth.

### Tissue processing and imaging

Animals were transcardially perfused with 0.9% saline and 4% paraformaldehyde solutions in 1X-PBS (15710-Electron Microscopy Sciences, Hatfield, PA). Brains and spinal cords were dissected and fixed overnight in 4% paraformaldehyde at 4°C. Spinal cords were embedded in 12% gelatin in 1X-PBS (G2500-Sigma Aldrich, St. Louis, MO), and 100µm transverse sections of the cervical spinal cord cut by Vibratome. A complete series of 120µm transverse sections were gathered from each brain. Sections were mounted on slides and imaged with a confocal microscope (Nikon AR1+) using 10X Plan Apochromatic (MRD00105, NA 0.45), 20X Plan Apochromatic (MRD00205, NA 0,75), or 60X Apochromatic Oil DIC N2 (MRD71600, NA 1.4) objectives. The emitted fluorescence from the sections was strong enough to image without the need to amplify using immunofluorescence methods. The anatomical locations of fluorescent CTB-488 and tdTomato in supraspinal sections were matched to the nomenclature of anatomical structures according to the Allen Brain Atlas. Cells labeled by CTB-488 and/or tdTomato were manually quantified using the taxonomy tool in NIS-Elements software.

### Immunohistochemistry

For 5-HT immunohistochemistry, brain stem sections containing raphespinal neurons were blocked in 10% goat serum containing 0.04% Triton-X, and incubated at room temperature for 1 hour while shaking. Brain stem sections were then transferred to a solution (3% goat serum, 0.04% Triton-X) containing the primary antibody, rabbit anti-5-HT (Sigma, S5545) at 1:1000 incubated for 18 hours at 4°C. Sections were then incubated for 2 hours at room temperature in secondary antibody, donkey anti-goat with a fluorescent tag, Alexa Fluor-647 (1:500) diluted in (3% goat serum, 0.04% Triton-X). To visualize glial fibrillary protein (GFAP), spinal sections were blocked as above and incubated with primary antibody overnight at 4 degrees (Z0334-DAKO, California, 1:500 RRID:AB10013482), rinsed, and then incubated for two hours with appropriate Alexa Fluor-conjugated secondary antibodies (Thermofisher, Waltham, MA, 1:500.). Following incubations, sections were mounted on glass slides and coverslipped with mounting medium. Sections were mounted on slides and imaged using a confocal microscope (Nikon AR1+) as above.

### DREADD activation and behavioral testing

Clozapine (Sigma, C6305) and was dissolved in 0.1M HCl too 1mg/ml, and diluted in saline to a final concentration of 1, 0.5, or 0.1µg/ml, then injected IP to mice to achieve final doses of 0.01, 0.05, 0.1, and 0.2 mg/kg (Gomez et al., 2017). CNO (Tocris, 9A/200456) was dissolved in saline to a concentration of 1µg/ml and injected IP to a final dose of 10mg/kg. Immediately after injection mice were placed in a clear-bottomed cage and digitally recorded from below. At 0min, 5min, 10min, 15 min, and then hourly thereafter, animal forelimb movement was scored on the following scale: 5=normal locomotion with predominantly plantar placement, 4 = weight-bearing locomotion but with frequent non-plantar placement (curled forelimb placed on ventral surface), 3=weight-bearing locomotion but entirely non-plantar placement, 2=no weight-bearing steps, but forelimbs make sweeping motions for forward propulsion 1=no weight-bearing steps, forelimbs make small movements that don’t move body, 0=forelimbs pinned with no discernible movement. In addition, transgenic CAMKIIa-Cre mice (Jackson labs, B6.Cg-Tg(Camk2a-cre)T29-1Stl/J) were tested on modified horizontal ladder task in which animals walked atop a wheel with variably spaced rungs (average 1.0cm gap, range 0.5 to 1.5cm). Animals were confined to the apex of the wheel, which rotated beneath them at a fixed speed of 0.78m/min. After one full rotation, data were collected for the next three rotations. Each right forelimb step was scored as correct (step centered on rung, which is grasped), wrist or toe placement errors (forelimb takes weight but is placed too far forward or back, such that rung is not grasped), or complete miss (wrist breaks the plane of the rungs). Animals were pre-trained on the task, and then tested in the absence of clozapine administration or 15 minutes after IP clozapine injection (0.1mg/kg).

### Tissue clearing and imaging

Animals were transcardially perfused with 0.9% saline and 4% paraformaldehyde solutions in 1X-PBS (15710-Electron Microscopy Sciences, Hatfield, PA). Whole Brains and spinal cords were dissected and fixed overnight in 4% paraformaldehyde at 4°C and washed 3 times in PBS pH 7.4 and stored in PBS. The dura was carefully and completely removed as residual dura can trap bubbles that prevent effective light-sheet microscopy. Samples were incubated on a shaker at room temperature in 50%, 80%, and 100% peroxide-free THF (Sigma 401757) for 12 hours each. Peroxides were removed from THF by passing 100% THF through a chromatography column filled with basic activated aluminum oxide (Sigma 199443) as previously described (Ertürk et al., 2012; Soderblom et al., 2015). The next day, samples were transferred to BABB solution (1:2 ratio of benzyl alcohol, Sigma, 305197; and benzyl benzoate, Sigma, B6630) for at least 3 h. After clearing, samples were imaged within 48 hours by light-sheet microscopy (Ultramicroscope, LaVision BioTec). The ultramicroscope uses a fluorescence macro zoom microscope (Olympus MVX10) with a 2× Plan Apochromatic zoom objective (NA 0.50). Image analysis and 3D reconstructions were performed using Imaris v8.4.2 software (Bitplane) after removing autofluoresence using the Imaris Background Subtraction function with the default filter width so that only broad intensity variations were eliminated. Artifact and nonspecific fluorescence surrounding the brain was segmented and removed using the automatic isosurface creation wizard based upon absolute intensity. Some of the cleared brains were also sliced to generate thick sections of 500-1000 microns and imaged on a FV1000 (Olympus) laser scanning microscope. Movies were generated using Imaris and merged in Photoshop CS6.

### Experimental Design and Statistical Analysis

An initial *in vivo* time course experiment was initiated with 15 adult female mice, sacrificed in equal groups at 3, 7, and 14 days post-injection. One animal in the 3 day group was lost to mortality, and one animal in the 7 day group was excluded on the basis of a mistargeted injection (no CTB-488 label detected in the spinal cord). The *in vivo* spinal injury experiment was initiated with five adult female mice in each group (injured versus uninjured). One injured animal was lost to mortality and one uninjured animal was excluded on the basis of mistargeted injections, leaving four animals in each group. To compare viral efficiency across all populations, the percent of dual CTB/viral labeling was quantified from at least 100 cells in each animal, and then compared across groups by two way ANOVA with post-hoc Tukey’s. To compare injured and uninjured animals, ANOVA with post-hoc Sidak’s tested for differences in transduction efficiency. For tissue clearing, four animals received cervical injection of Retro-AAV-tdTomato to cervical spinal cord and were sacrificed 14 days later, and three adult female mice received both cervical AAV2-Retro-EGFP and lumbar AAV2-Retro-tdTomato and were sacrificed four weeks later. DREADD experiments were initiated with 5 wildtype and 5 CAMKII-Cre female mice injected with AAV2-Retro-DREADD constructs, and three uninjected wildtype females. One Camkii-Cre was lost to mortality. Slip rates in the presence or absence of clozapine stimulation on the ladder task were compared by paired t-tests.

## RESULTS

### AAV2-Retro effectively transduces reticulospinal, rubrospinal, and corticospinal neurons

AAV serotypes vary widely in their tropism for different types of neurons and their onset of transgene expression (Castle et al., 2016). This appears to be particularly true for retrograde transduction from the spinal cord, as prior tests showed higher efficiency in propriospinal and brainstem populations compared to CST neurons (Jara et al., 2012; Klaw et al., 2013). To comprehensively profile the efficiency and time-course of retrograde transduction by AAV2-Retro in supraspinal populations, fifteen adult female mice were injected with a mixture of the retrograde tracer CTB-488 and AAV2-Retro-tdTomato, bilaterally to cervical spinal cord. Animals were sacrificed at 3, 7, or 14 days post-injection and complete series of 120µm sections of the brain and cervical spinal cord was collected. CTB-488 is a widely used retrograde tracer that efficiently labels supraspinal neurons and was used as a positive control to mark neurons that projected axons to the site of injection. Sections were imaged by confocal microscopy and transduction efficiency was quantified by the percent of CTB-488+ neurons in each population that co-expressed tdTomato. A summary and statistical comparison of transduction efficiency across all populations and timepoints is provided in **Supplementary Figure 1**, and detailed description of selected nuclei is provided below.

*Injection site and cervical propriospinal neurons* – Spinal cord sections were collected from the site of injection and at 1mm and 2mm rostral. Injection sites showed intense CTB-488 and tdTomato signal at all time points, confirming appropriate targeting and transduction of cell bodies directly exposed to AAV2-Retro (**Fig. 1B,C,D**). At locations 1mm and 2mm rostral to the site of injection, numerous cell bodies were labeled with CTB-488+, indicating retrograde transport of CTB-488 (**Fig. 1E-G**). Of these, fewer than 20% at 1mm and fewer than 10% at 2mm co-expressed tdTomato at any time point (**Fig1. H-N**; 1mm: 10%, 14% and 19% at 3,7, and 14 days; 2mm 3%, 3%, and 6% at 3,7, and 14 days). Interestingly, despite the low transduction in spinal neurons, by 14 days tdTomato label was detected in white matter locations corresponding to descending tracts, including the ventral aspect of the dorsal column, the location of CST axons (arrow, **Fig. 1M**). This axonal labeling hinted at more effective transduction in higher supraspinal populations, which we confirm below. These data show successful targeting of injections and transduction at sites of spinal injection, but indicate relatively low retrograde transduction efficiency in nearby propriospinal populations in upper cervical spinal cord.

**Figure 1:**
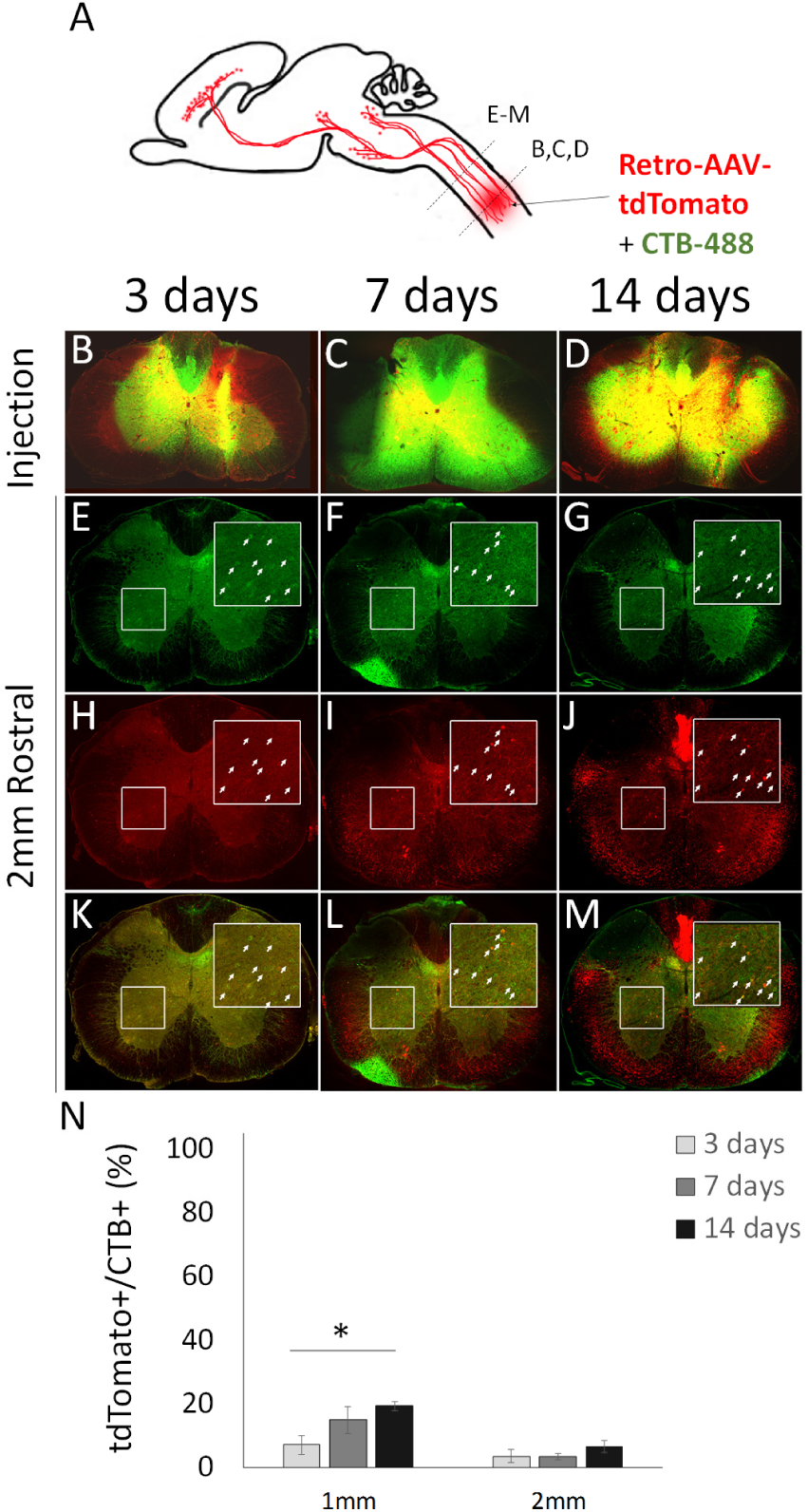
AAV2-Retro shows limited retrograde transduction of cervical propriospinal neurons. (A) Mixed AAV2-Retro-tdTomato and CTB-488 were injected bilaterally to C4/5 grey matter, and transverse spinal sections at the injury site and 1mm or 2mm rostral were examined 3, 7, or 14 days later. (B-D) Transverse sections at the level of spinal injection show readily detectable CTB-488 and tdTomato. (E-G) In cervical spinal cord 2mm rostral to the injection, CTB-488 retrogradely labels cell bodies that project axons to the injection site (arrows). (H-M) tdTomato signal is rarely detectable in CTB-488+ cell bodies, indicating low viral expression. (N) Quantification of tdTomato detection in CTB-488+ cell bodies shows fewer than 20% of propriospinal neurons projecting to C4/5 were transduced 1mm rostral to the injury, and fewer than 10% at 2mm. * p<.05, 2-way ANOVA with post-hoc Tukey’s. Scale bar is 1mm. N>100 individual cells from each of at least four animals at each time point. Scale bar is 1mm.

*Brainstem and Cerebellar neurons:* Spinal projection neurons throughout the brainstem were labeled by CTB-488. Based on anterior/posterior position in the sequence of sections and nearby neuroanatomical landmarks, we identified clusters of CTB-488 positive cells corresponding to reticulospinal, cerebellospinal, and vestibulospinal nuclei (**Fig. 2A-G**). By three days post-injection, AAV2-Retro-tdTomato signal was readily detectable in a majority of these cells (reticulospinal: 61.2 (±7.4SEM)%, 88.6 (±1.6SEM)%, and 67.4 (±5.8SEM)%, cerebellospinal: 98.6 (±1.4SEM)%, 100 (±0SEM)%, and 88.8 (±5.2SEM)%, vestibulospinal: 80.8 (±1.4SEM)%, 96.5 (±0.9SEM)%, and 68.5 (±6.4SEM)%, at 3, 7, and 14 days post injection, respectively (**Fig. 2H**). Next, using 5HT immunohistochemistry, we identified raphe neurons located near the medullary ventral midline (**Fig. 2I-L**). In striking contrast to other brainstem-spinal populations, tdTomato signal was absent from 5-HT-positive raphe neurons at all time points, including cells in which CTB-488 signal that confirmed axonal projection to the site of spinal injection (**arrows, Fig 2L**). These data indicate cell-type tropism that prevents effective transduction of serotonergic neurons. Consistent with this, the initial characterization of AAV2-Retro indicated preferential retrograde transduction of glutamatergic neurons and limited transduction of dopaminergic neurons (Tervo et al., 2016). In summary, these results indicate that AAV2-Retro effectively and rapidly transduces most brainstem-spinal populations, with the notable exception of raphe-spinal projection.

**Figure 2:**
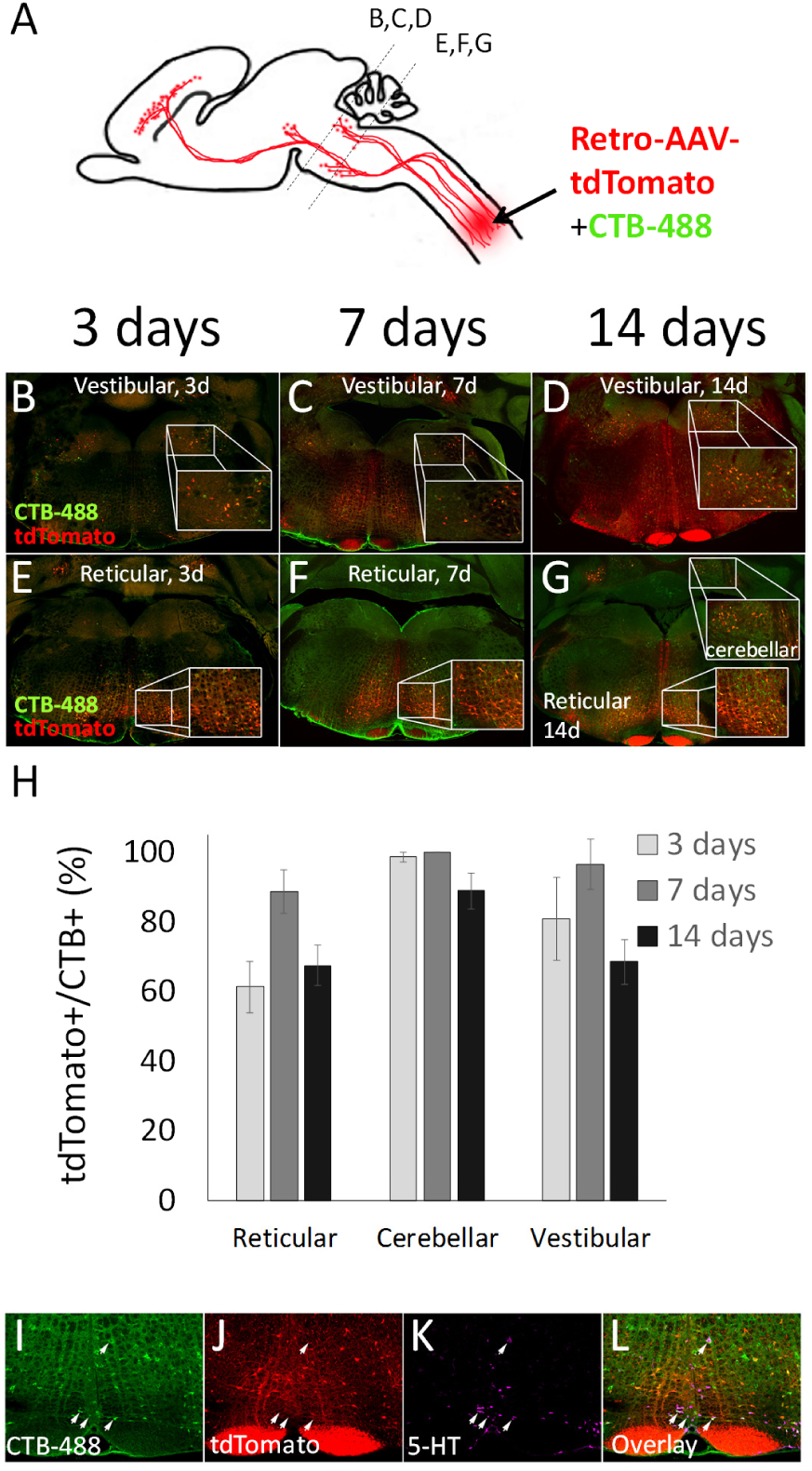
AAV2-Retro effectively transduces brainstem-spinal projection neurons (A) Mixed AAV2-Retro-tdTomato and CTB-488 were injected bilaterally to C4/5 grey matter, and transverse sections of brainstem examined 3, 7, or 14 days later. In vestibulospinal (B-D), reticulospinal (E-G), and cerebellar-spinal (E-G) populations, spinally projecting neurons identified by CTB-488 also express tdTomato as early as three days post-injection. (H) Quantification of the percent of CTB-488+ neurons that are virally transduced shows a majority of vestibulospinal, reticulospinal, and cerebellar-spinal neurons are retrogradely labeled. N>100 individual cells from each of at least four animals at each time point. Groups did not differ significantly (p>0.05, 2 way ANOVA with post-hoc Tukey’s). (I-L) Two weeks after spinal injection of mixed CTB-488 and AAV2-Retro-tdTomato, transverse brainstem sections were immunostained for 5HT. Raphe-spinal neurons (arrows) were identified by CTB-488 (I) to confirm an axon projection to the C4/5 injection site, and expression of 5HT (K). These neurons were invariably negative for tdTomato, although nearby non-serotonergic neurons were effectively transduced.

*Red Nucleus*- The red nucleus spans the boundary of the diencephalon and midbrain and is composed of two distinct populations of neurons, the rostral parvocellular, and the caudal magnocellular. Both populations project axons to the spinal cord and, as expected, were effectively labeled by retrograde CTB-488 (**Fig. 3A-D, K-M**) (Liang et al., 2012a). In the magnocellular nucleus virtually all spinally projecting neurons were transduced, with 97.5 (±1.5SEM)%, 100 (±0.7SEM)%, and 99.3 (±0.5SEM)% co-labeled at 3,7,and 14 days post injection (**Fig. 3E-J,T)**. In the parvocellular division, 70.9 (±8.0SEM)%, 93.3 (±4.9SEM)%, and 82.9 (±2.0SEM)% of CTB-488+ cells expressed tdTomato at 3, 7, and 14 days post injection, respectively (**Fig. 3N-T)**. Thus spinally injected AAV2-Retro effectively transduces cell bodies in the red nucleus.

**Figure 3.**
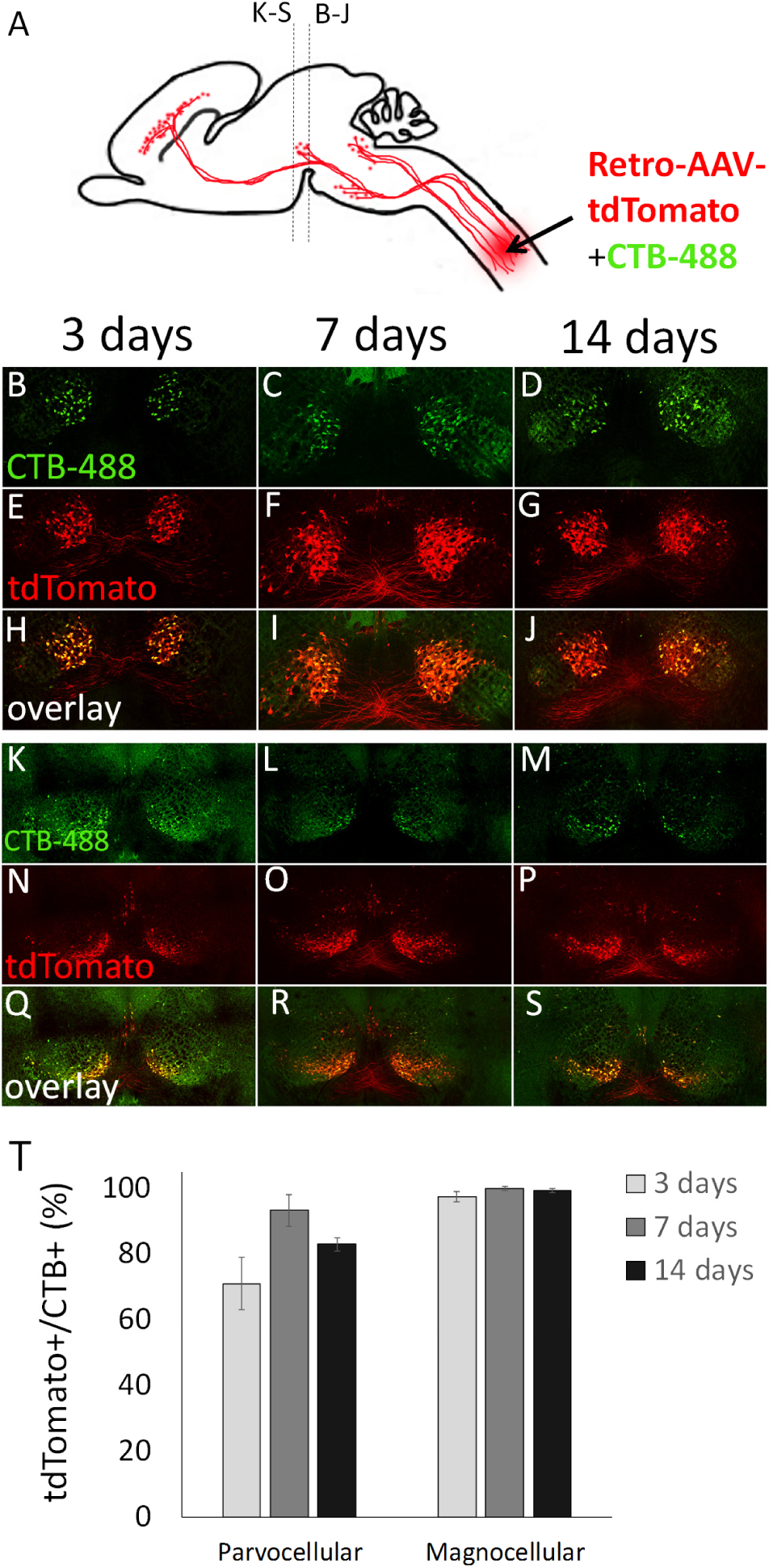
Effective retrograde transduction of rubrospinal neurons. (A) Mixed CTB-488 and AAV2-Retro-tdTomato were injected to cervical spinal cord and transverse sections of midbrain prepared 3, 7, or 14 days later. (B-J) show the magnocellular population, (K-S) the parvocellular (T) A majority of CTB-488+ neurons in both parvo- and magnocellular nuclei were co-labeled with tdTomato, indicated rapid and effective transduction. Groups did not differ significantly (p>0.05, 2 way ANOVA with post-hoc Tukey’s). N>100 individual cells from each of at least four animals at each time point.

*Corticospinal*. CST neurons reside in layer V of sensorimotor cortex and provide the only direct input from cortex to spinal cord. Until recently, retrograde vectors have displayed a limited ability to transduce CST neurons. We examined transverse cortical sections between 3mm anterior and 2mm posterior to Bregma, which spans all regions of cortex that project axons to the spinal cord (for a more complete description of distinct CST populations see 3D visualization results below). As expected, retrograde CTB-488 was readily detected in CST cell bodies at all time points (**Fig. 4A-E**). Virally expressed tdTomato was expressed in more than 80% of CTB-488+ cell bodies by just 3 days post-injection, and in more than 95% of CTB-488+ cell bodies by 7 and 14 days (**Fig 4F-N**). As noted earlier, tdTomato signal emerged in CST projection tracts in the brainstem by 7 days post-injection (**Fig. 2C,F**), and in the spinal CST tract by 14 days (**Fig. 1J,M**). These data indicate that cervical injection of AAV2-Retro results in expression in most CST neurons within three days, and that within two weeks fluorophore expression and transport are sufficient to detect the projecting axons in cervical spinal cord.

**Figure 4.**
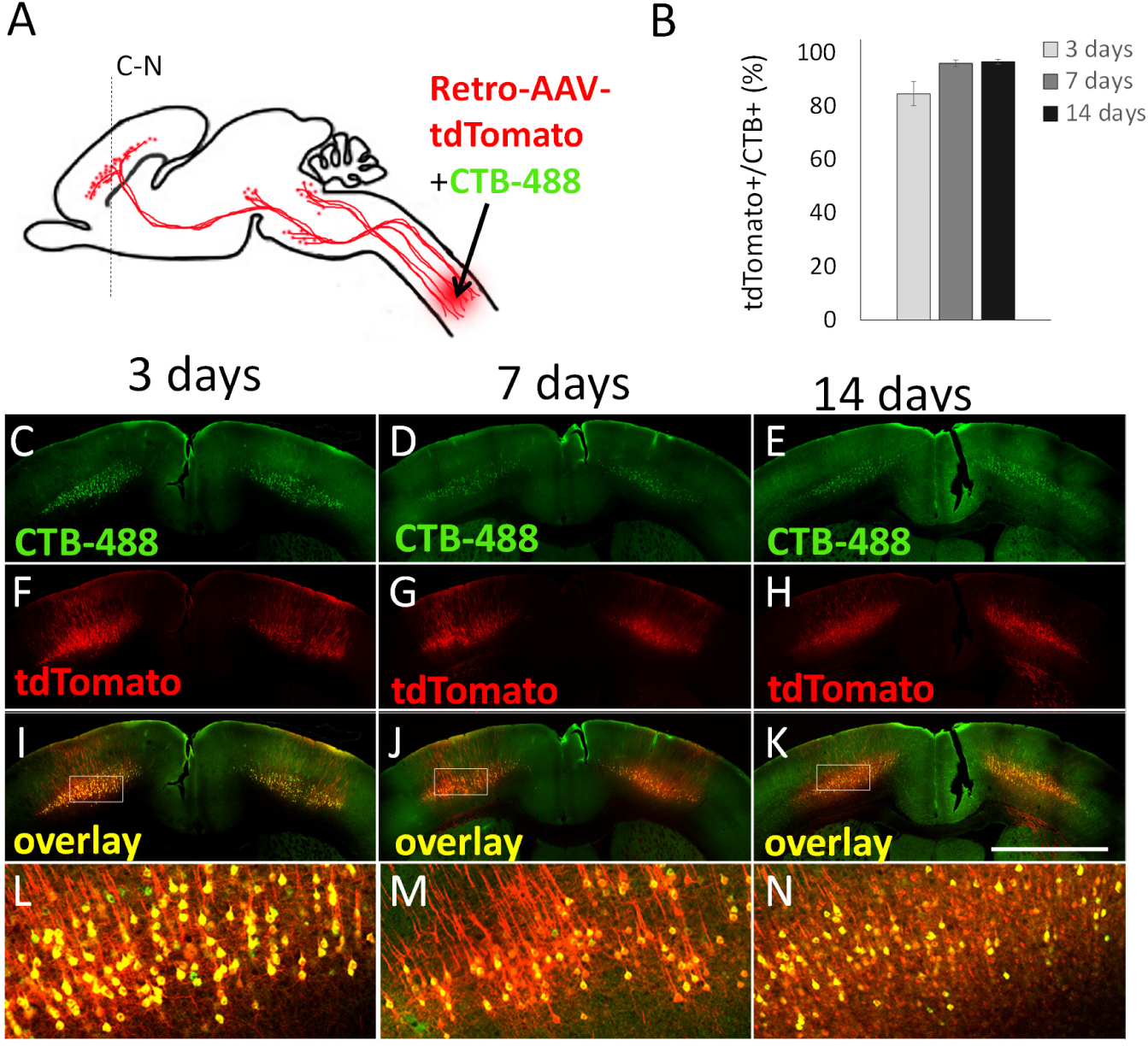
Effective retrograde transduction of corticospinal tract neurons. (A,ES) Adult mice received cervical injection of mixed AAV2-Retro-tdTomato and CTB-488, and the percent of double-labeled cells quantified in transverse sections of cortex 3, 7, or 14 days later. (C-E) CTB-488 identifies CST neurons. (F-H) tdTomato is readily detectable as early as 3 days post-injection). (I-N) Overlays show the great majority of CTB-488+ cells also express tdTomato, indicating efficient retrograde transduction. Groups did not differ significantly (p>.05, ANOVA with post-hoc Sidak’s). N≥4 animals per time point, >100 cells/section, three sections per animal.

### Retrograde transduction remains effective following cervical spinal injury

A pre-requisite for potential therapeutic utility is that retrograde vectors display effective retrograde transduction when applied to axons after injury. We therefore tested AAV2-Retro administration after cervical spinal injury. Four adult female mice received unilateral dorsal hemisections of C4/5 spinal cord, which completely severs the dorsal and dorsal-lateral CST axons on the right side of the spinal cord. Immediately after injury, animals received bilateral injections of mixed AAV2-Retro-tdTomato and CTB-488 to C4 spinal cord, 0.5mm rostral to the injury. As control, four additional animals received AAV2-Retro-tdTomato/CTB-488 injections without injury. Animals were sacrificed at 28 days post-injection and a complete series of 120µm sections of the brain was collected, along with sagittal sections of the spinal injury site. Immunohistochemistry for GFAP at the site of spina injury confirmed lesion depth in all animals, and in the dorsal cord tdTomato and CTB-488 signal was confined to tissue rostral to the injury, indicating complete ablation of dorsal axon tracts (**Fig. 5A-E**). As expected from this partial injury, tdTomato+ axons continued uninterrupted in ventral tracts, indicating likely sparing of other supraspinal neurons that project axons ventrally. We therefore focused our analysis on the CST neurons, a population unequivocally affected by dorsal injury. As expected, in uninjured animals CTB-488 retrogradely labeled cortical cell bodies, and the great majority of these expressed tdTomato (right cortex: 93.96(±2.9SEM)%, left cortex 93.9 (±2.0SEM)%) (**Fig. 5F,H**). Similarly, in the right (spared) cortex of animals that received unilateral spinal transection, transduction efficiency was 90.3 (±5.7%) (**Fig. 5F, H**). In the left (injured) cortex, efficiency was 98.8 (±1.1%), not significantly different from either contralateral cortex or uninjured animals (p<.05, ANOVA with post-hoc Sidak’s) (**Fig. 5G, H**). Consistently, the absolute numbers of tdTomato+ cells counted in five tissue sections was similar in right and left cortex (right: 339.3 (±71.7SEM), left: 400.7 (±42.0SEM), p=0.51, paired t-test). Thus AAV2-Retro-tdTomato is equally effective in transducing CST neurons when applied acutely after a spinal injury.

**Figure 5.**
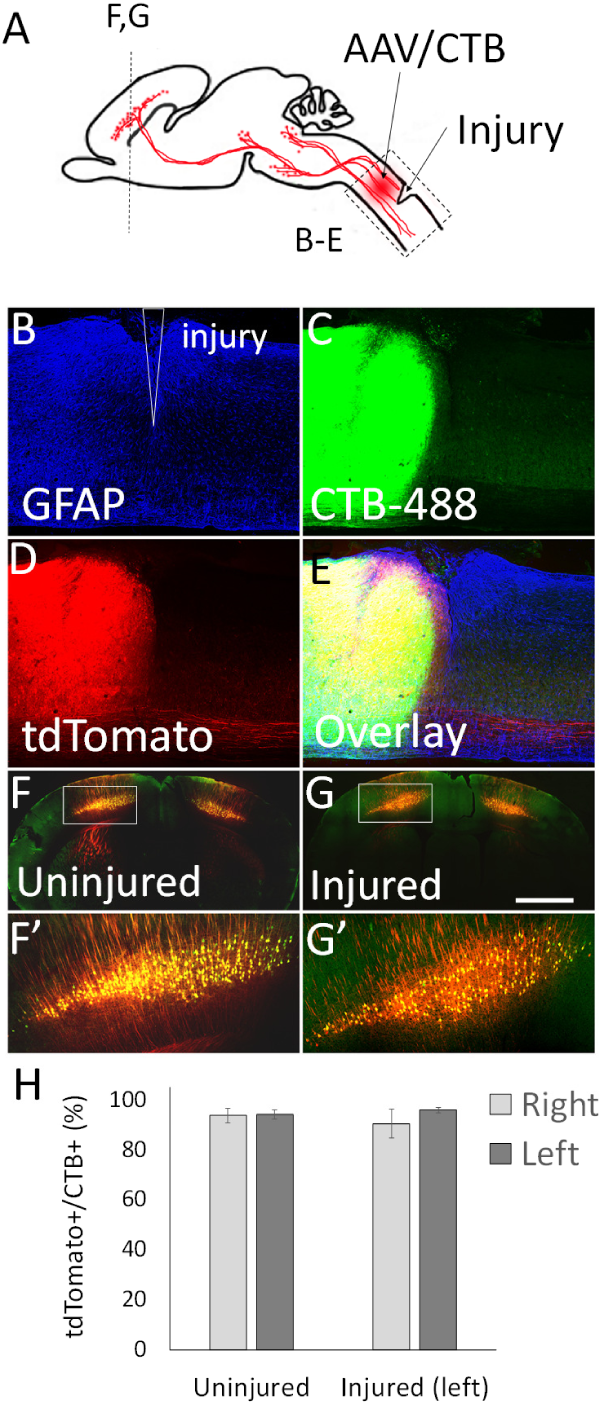
Effective retrograde transduction of corticospinal tract neurons after spinal injury. (A) Adult female mice received unilateral cervical dorsal hemisection or sham injury, followed by bilateral injection of mixed AAV2-Retro-tdTomato and CTES-488. (B-E) In sagittal sections of spinal cord 28 days post-injection, GFAP immunohistochemistry (B) and the absence of CTB-488 (C) and tdTomato (D) signal caudal to the injury confirmed complete dorsal transections. (F-G) Co-expression of CTB-488 and tdTomato was high in CST neurons in injured animals (F) and in both cortices of animals that received unilateral spinal injury (G). (H) Quantification of co-expression rates showed no difference in the left cortex of injured animals, the location of axotomized CST cell bodies (All 2-way comparisons p>.05, 2-way ANOVA with post-hoc Tukey’s). N=four animals per group, >200 individual cells quantified per animal.

### Tissue Clearing and 3D reconstruction reveals spatial organization of descending spinal inputs

Advances in tissue clearing and 3D microscopy now enable the visualization of neurons and their projections in intact tissue. We therefore injected AAV2-Retro-tdTomato bilaterally to cervical spinal cord, and then two weeks later performed tissue clearing and 3D-imaging of brain tissue (Bray et al., 2017). Consistent with the section-based histology discussed above, 3D imaging of intact tissue revealed widespread fluorophore expression in supraspinal nuclei in the brain and in descending axon tracts. In addition, the 3D imaging provided new insights by revealing anatomical features of supraspinal input that were difficult to detect in the 2D tissue sections.

First, 3D imaging clarified the distribution of corticospinal tract neurons into three distinct groups: a large central mass corresponding to primary sensorimotor cortex, a distinct group located rostrally, corresponding to secondary motor cortex or rostral forelimb area (RFA), and a caudal/lateral group corresponding to secondary sensory cortex (**Fig. 6A-C, and Supp. Movie 1**). Axons from each area could be clearly seen descending from various locations, first forming the corona radiata and then converging into the internal capsule. Axons from the caudal/lateral S2 group converged with the main CST tract just above the cerebral peduncle, and all CSTs descended as a unified tract at that point (**Fig. 6A-C**). Descending CSTs could be distinguished as they descended into the pyramids and across the medullary decussation. Interestingly, the 3D reconstruction revealed clear points of collateralization along this path (**Fig. 6A-D, arrows; Supp. Movies 1-4**). First a cloud-like structure emanated just below the corona radiata and projected into the putamen (**Fig. 6A-D**, **purple arrow**). Second, extending dorsally from the cerebral peduncles was a tract destined for the pre-tectal and tectal areas (**Fig. 6A-D, orange arrows; Supp. Movie 3**). In addition, near the rostral end of the pyramidal tract, a dense projection extended ventrally, likely corresponding to collateral innervation of the basilar pontine nuclei ((**Fig. 6A-D, green arrows; Supp. Movies 1-3**). We emphasize that only axons that reached cervical spinal cord were labeled, and thus these collateral tracts clearly indicate the presence of branched projections from definitive CST neurons. To corroborate this finding we examined tracings of individual neurons recently available at the Mouselight Neuron Browser (https://www.janelia.org/project-team/mouselight) and identified six cortical neurons with unequivocal projection to the spinal cord (Economo et al., 2016). Interestingly, all six individual neurons showed branching into pre-tectal areas, five branched into basilar pontine nuclei, and two branched into the putamen (**Fig. 6E**). Thus, CST axons collateralize extensively in supraspinal nuclei.

**Figure 6.**
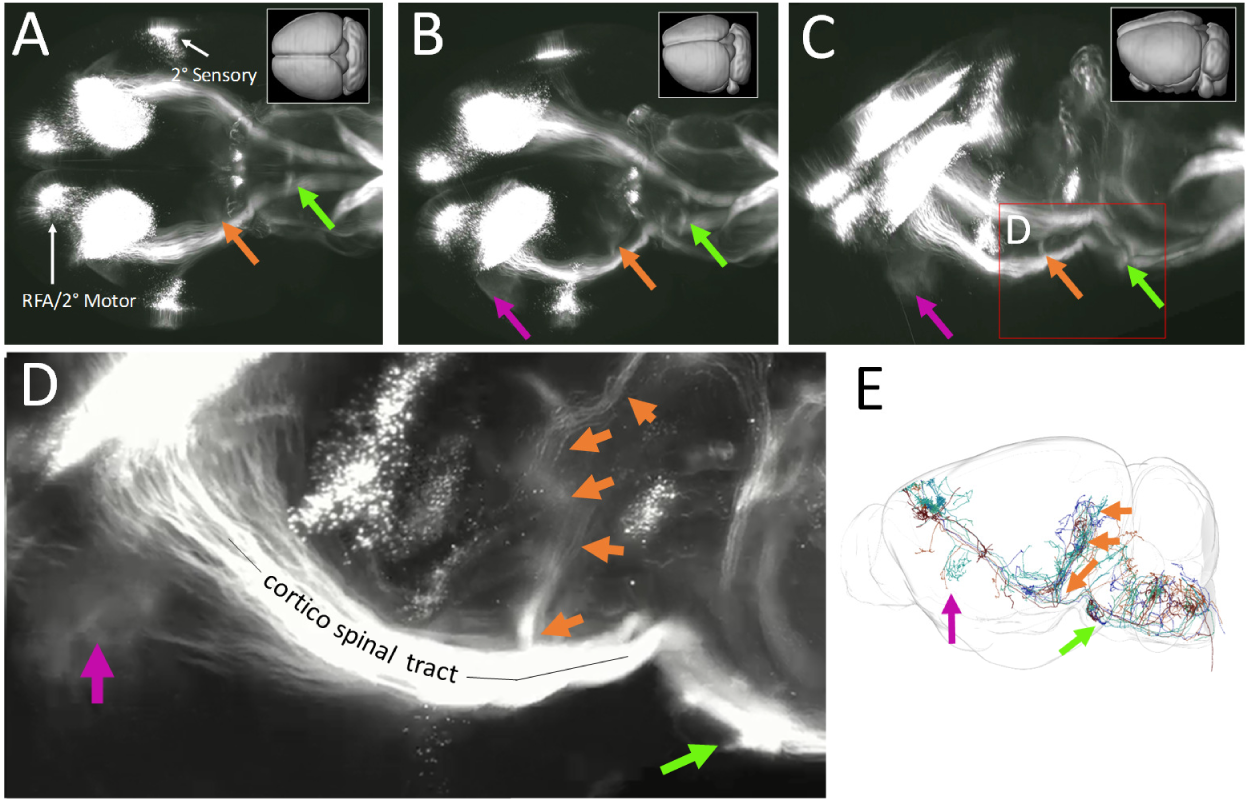
Retro-AAV and 3D imaging reveal CST cell body distribution, axon trajectories, and collatoralization. (A-D) Adult mice received bilateral cervical injection of AAV2-Retro-tdTomato. Two weeks later brains were optically cleared by 3DISCO and imaged with light sheet microscopy Images showed rotated views of whole brain, with rostral to the left and initially viewed from the dorsal surface in A. Three distinct groups of CST cell bodies are apparent, and to clear points of collateral branching from the main CST are visible (arrows). (E) Individual tracings of CST neurons from the Mouselight Neuron Browser. Consistent with the 3D reconstruction, collateral branches to tectal areas (orange arrow) and to Basilar Pontine Nuclei (green arrow) are visible.

3D imaging of the intact brain also clarifies the spatial relationship of supraspinal nuclei and can help draw attention to less-studied populations. For example, 3D imaging of the forebrain reveals not only the CST populations described above, but also supraspinal populations located in the ventral forebrain. These are likely hypothalamic-spinal populations (Liang et al., 2011), appearing here as scattered cells in the lateral and paraventricular hypothalamus (**Fig. 7A, B, Supp. Movie 4)**. Similarly, 3D imaging of the red nucleus showed the relative position of the parvocellular and magnocellular populations and the differences in cell morphology between them (**Fig 7C-E, Supp. Movie 5**). Also apparent in the 3D imaging is a prominent midline group of cells corresponding to the centrally projecting Edinger Westphall nucleus (EWcp) as well as more diffusely distributed cell bodies of the interstitial nucleus of Cajal (Kozicz et al., 2011; Liang et al., 2011) (**7C; Supp. Movie 5**).

**Figure 7.**
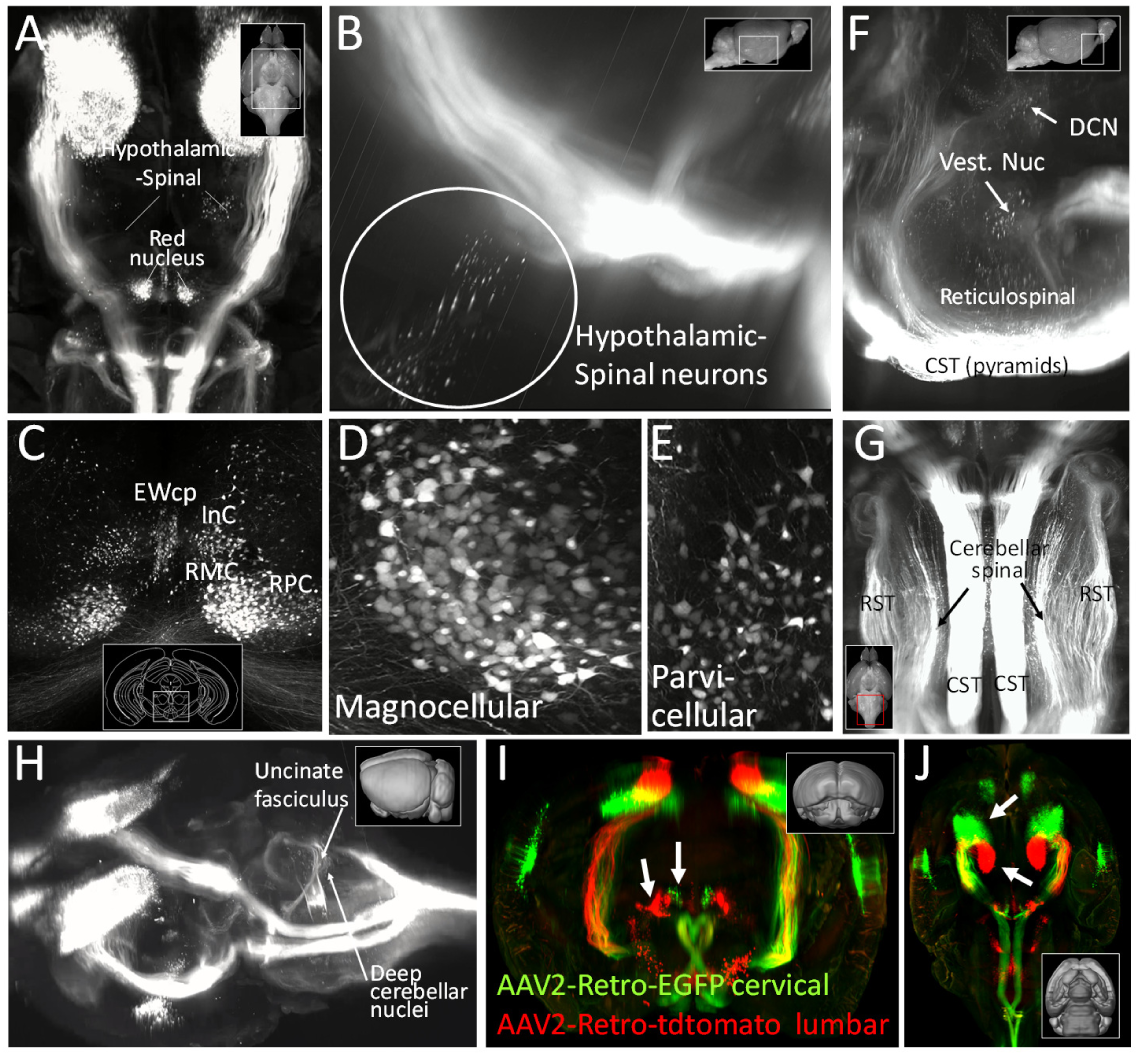
Retro-AAV and 3D imaging reveal supraspinal populations and axon trajectories. **(A-G)** Adult mice received bilateral cervical injection of AAV2-Retro-tdTomato, and two weeks later brains were optically cleared by 3DISCO and imaged with light sheet microscopy. **(A,B)** Imaging in the diencephalon shows scattered cell bodies likely corresponding to previously described hypothalamic-spinal populations. **(C-E)** 3D imaging in the midbrain shows magnocellular (RMC) and parvocellular (PRC) populations of the red nucleus, as well as the centrally projecting component of the Edinger Westphal nucleus (Ewcp) and the insterstitial nucleus of Cajal (INC). **(F,G,H)** 3D imaging shows multiple brainstem populations and axon tracts, including the DCN (dorsal cerebellar nuclei), vestibular nuclei, reticulospinal populations, the corticospinal tract (CST), rubrospinal tract (RST) and cerebellospinal tract. **(I,J)**. Adult mice received bilateral injections of Retro-AAV-EGFP to cervical spinal cord, and Retro-AAV-tdTomato to lumbar spinal cord. Brains were cleared and imaged four weeks later. Rubrospinal neuronal cell (arrows, I) and CST cell bodies (arrows, J) segregated into distinct lumbar vs. cervically-projecting populations.

In the cerebellum and brainstem, 3D imaging also helps resolve some of the complexity of the multiple supraspinal populations and their various axon tracts, including spinally projecting reticular neurons, vestibular neurons, and deep cerebellar nuclei **(Fig. 7F, Supp. Movie 6)**. Also apparent in the brainstem are the details of trajectories of descending axons. For example, rubrospinal axons enter the upper brainstem as a confined bundle, separate in into discrete bundles in the lower pons and upper medulla, then re-converge prior to entering the spinal cord (**Fig. 7G; Supp. Movie 6)**. The trajectory of the cerebellar-spinal tract could also be discerned, first extending to the contralateral cerebellum via the uncinate fasciculus (**Fig. 7H, arrow**), then through the superior cerebellar peduncle, and then descending in a position just lateral to the rubrospinal tract. Similar to the rubrospinal tract, the cerebellar-spinal tract broke into discrete bundles as it descended through the brainstem, and then reaggregated just before entering the spinal cord. (**Fig. 7G**; **Supp. Movie 6**). Overall, combining retrograde fluorescent labeling with 3D tissue reconstruction offers a clearer view of the anatomy of supraspinal projections.

Finally, we used combined AAV2-Retro and 3D imaging to visualize the relative positions of neurons that project to cervical versus lumbar spinal cord. AAV2-Retro-EGFP was injected bilaterally to cervical spinal cord of adult mice, and AAV2-Retro-tdTomato to lumbar spinal cord. In pilot studies we discovered that unlike cervical injections, lumbar injection of AAV2-Retro required a full four weeks to achieve adequate expression level, and accordingly survival times were increased from two weeks to four. Consistent with prior work, lumbar-projecting CST neurons (tdTomato+) were confined to cortex located just caudal to the main population of cervically projecting CST neurons (EGFP+) (Kamiyama et al., 2015) (**Fig. 7I,J, Supp. Movie 7**). We did not observe any CST neurons that were double-labeled, and no tdTomato+ neurons were located in secondary sensory cortex or in the rostral forelimb area, emphasizing the presence of neurons with projections to only lumbar or cervical cord. Similarly, in the red nucleus, cervically-versus lumbar-labeled neurons segregated into two discrete populations, with the lumbar-projecting neurons located lateral to the cervical population with very few double labelled neurons (Flumerfelt and Gwyn, 1974; Liang et al., 2012a) (**Fig. 7J, Supp. Movie 7**). Thus, 3D imaging can also assist in the visualization of the topographical distribution of cell bodies within supraspinal populations that extend axons to different positions along the rostal-caudal axis of the cord. Overall, 3D reconstruction confirms the wide distribution of supraspinal neurons transduced by AAV2-Retro.

### Retrograde DREADD silencing of supraspinal populations

The histological data indicated successful gene delivery to the majority of supraspinal populations with input to cervical spinal cord. To test the functional consequences of silencing the affected neurons we utilized a DREADD-based strategy, shown previously to effectively silence a variety of supraspinal inputs (Hilton et al., 2016; Siegel et al., 2015; Wahl et al., 2014). We used a Cre-dependent Flex-DREADD construct, in anticipation of future work involving more controlled expression, but in initial experiments deployed the construct in a manner that aimed for widespread and non-specific DREADD expression. We co-injected AAV2-Retro-Flex-DREADD along with AAV2-Retro-Cre to cervical spinal cord of adult mice, such that all supraspinal neurons that project to cervical spinal cord would have access to both DREADD and the activating CRE. Indeed, mCherry fluorescence confirmed robust DREADD expression in brainstem, red nucleus, and cortical neurons (**Fig. 8A**).

**Figure 8.**
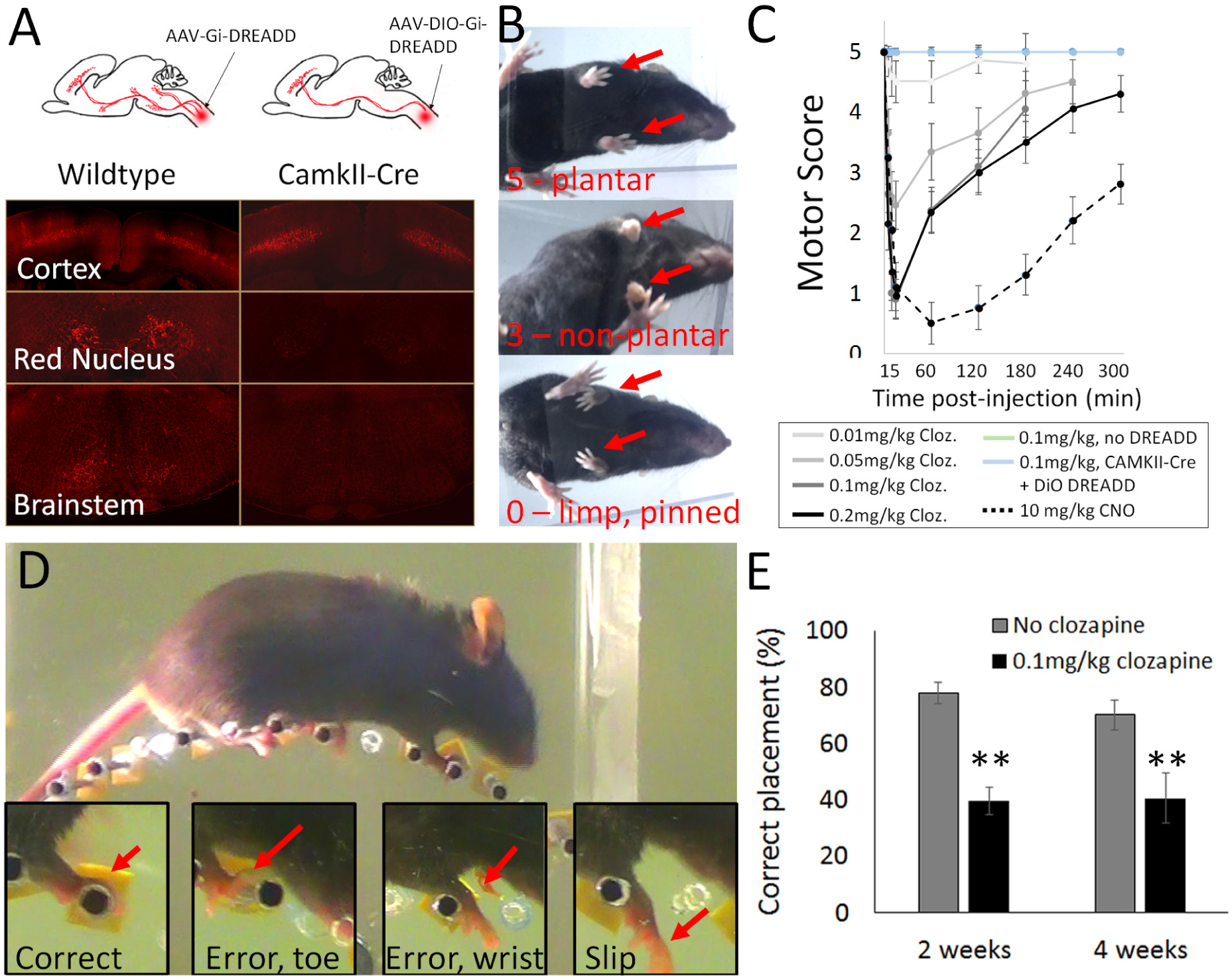
Gi-DREADD mediated silencing of supraspinal input to cervical spinal cord. (A) Mixed AAN/2-Retro-Flex-Gi-DREADD-mCherry and AAN/2-Retro-Cre was injected to the cervical spinal cord of wildtype mice, and AA2-Retro-Flex-Gi-DREADD-mCherry alone was injected to the cervical spinal cord of CAMKII-Cre animals. Four weeks later, mCherry signal was readily detectable in the brainstem, red nucleus, and cortex of wildtype animals, but was cortically enriched in Camkll-Cre animals. (B) Clozapine produce reproducible forelimb deficits in DREADD-injected wildtype animals. (C) Dose response and timing curves for clozapine and CNO-triggered motor deficits. Both ligands caused reversible forelimb paralysis in Gi-DREADD injected wildtype animals, but not Camkll-Cre animals or non-injected controls. (D) Mice were tested on a horizontal ladder task atop a wheel with irregularly spaced rungs and errors in forelimb placement were scored. (E) In CAMKII-Cre animals, clozapine produced significant reduction in correctly targeted steps, consistent with selective silencing of corticospinal tract input to the spinal cord. N=5 wildtype, 4 Camkll-Cre, and 3 non-injected controls. **p<.01, paired t-test.

Starting two weeks after viral injection, we activated DREADD receptors using clozapine. shown by a recent report to function as a DREADD ligand (Gomez et al., 2017). We initially delivered clozapine IP at a concentration of 0.1mg/kg, a concentration reported to achieve effective DREADD activation, and observed animals in open field locomotion. Within 5 minutes, animals displayed clear changes in forelimb movement, with a disappearance of plantar placement and the emergence of weight-bearing steps on the curled dorsal surface (**Fig. 8B**). Within 15 minutes, weight bearing steps by forelimbs were absent and limbs were pinned beneath the torso. Careful examination showed small, non-effective movements is some animals, and no forelimb movement in others. By two hours after injection, most animals returned to weight-bearing steps with non-plantar placement, and by five hours showed near-normal locomotion with only occasional non-plantar placement (**Fig. 8B,C**). To quantify these effects, we established a simple five-point motor scale from 5 (normal forelimb stepping), 4 (occasional non-plantar steps) 3 (weight bearing steps but completely non-plantar), 2 (no weight bearing steps but occasional large sweeping movements for forward propulsion), 1 (limb predominantly immobile but occasional small movements) and 0 (no discernable movement). We administered a range of clozapine doses, taking advantage of the reversibility of the effects to test a different dose each day. At 0.01mg/kg, clozapine produced only a partial loss of plantar placement, and at 0.05mg/kg a complete loss of plantar placement but still some weight support. Clozapine at concentrations of 0.1mg/kg and 0.2mg/kg produced similar curves, with maximal impairment at 15 minutes post-injection (0.90 (±0.34SEM) and 0.95(±0.39SEM), respectively). These data establish a dose-response curve for clozapine-mediated DREADD activation, with maximal effects achieved at 0.1mg/kg. At all doses, effects were maximal at 15 minutes, partially recovered within an hour, and mostly reversed by five hours (**Fig. 8C**). Finally, we compared these effects to DREADD activation with clozapine N-oxide (CNO), an inert compound that has been widely used for DREADD studies, but which was recently reported to be converted to clozapine (Gomez et al., 2017). Consistent with this model, CNO produced paralysis similar to clozapine, but with a delayed and extended time course; effects were maximal at 1 hour and only partially recovered by five hours (**Fig. 8C**). Taken together, these data indicate that retrograde delivery of inhibitory DREADDs to cervical spinal cord affects a portion of descending motor input that is sufficient to abolish effective forelimb movement.

To render DREADD expression more specific to CST neurons we injected AAV2-Retro-Flex-Gi-DREADD-mCherry to the cervical spinal cord of CAMKIIa-Cre animals, in which Cre expression is highly enriched in glutamatergic projection neurons in the forebrain (Tsien et al., 1996). In this way, DREADD expression would occur predominantly in corticospinal tract neurons. Indeed, DREADD-injected CamkIIa-Cre animals showed limited expression of mCherry in brainstem and red nucleus but high expression layer five of cortex (**Fig. 8A**). In these animals clozapine injections produced no hint of the complete paralysis seen previously and no changes in plantar placement (**Fig. 8C**), consistent with prior literature showing CST to be dispensable for gross locomotion in mice (Starkey et al., 2005). To test for effects on skilled locomotion we employed a modified horizontal ladder task, in which mice were placed atop a wheel with irregularly spaced rungs, which was rotated beneath animals at a controlled and constant speed (**Fig. 8D**). The placement of forelimbs was scored by slow-motion video analysis. Clozapine (0.1mg/kg) produced a highly repeatable and reversible increase in errors in forelimb placement, which is consistent with selective silencing of CST input (**Fig. 8E)**. Combined, these data highlight the potential of AAV2-Retro-DREADD, combined with transgenic targeting strategies, to probe the functional contribution of individual supraspinal inputs.

## DISCUSSION

This work highlights new approaches for the study and manipulations of supraspinal input to the spinal cord. We showed that AAV2-Retro is broadly effective in transducing most supraspinal populations, including the corticospinal tract, and that transduction is unaffected by acute axon injury. When combined with tissue clearing and 3D reconstruction, retrograde delivery of fluorophores provides interesting insights into the anatomy of descending axon tracts. Finally, retrograde delivery of inhibitory DREADDs is sufficient to block forelimb movements, highlighting the potential for functional dissection of supraspinal inputs to the spinal cord. These data show an important advance in the capacity for retrograde gene delivery to act as a therapeutic tool, as well as to provide anatomical and functional insight into spinal pathways.

### AAV2-Retro tropism

The feature that most distinguishes AAV2-Retro from other AAV serotypes is its highly effective retrograde transduction of CST neurons. AAV1, -2, -5, -6, -7, -8, or -9 encoding GFP yields retrograde expression in 0 to 21.7% of CST neurons, even with antibody amplification (Jara et al., 2012). Similarly, AAV1, -2, -5, -8, or -9 delivered to the injured spinal cord drives retrograde transduction of brainstem and rubrospinal neurons but not CST (Klaw et al., 2013). The lentiviral HiRet vector was recently reported to achieve high retrograde transduction rates in CST neurons when injected to early postnatal animals, but these studies left unanswered whether HiRet is similarly effective in adults (Liu et al., 2017; Wang et al., 2017). Here we found that spinal injection of AAV2-Retro in adult animals transduces CST neurons with more than 95% efficiency. Fluorophore expression is quite strong, requires no antibody amplification, and can be visualized within days of injection. AAV2-Retro thus stands out as a powerful new tool for gene delivery to supraspinal populations, including CST.

AAV2-Retro was not as effective in labeling some supraspinal neurons. Although some brainstem nuclei (e.g.vestibular) were transduced at high rates, others showed lower efficiencies (**Fig. 2**). In particular, raphe-spinal neurons were not transduced by AAV2-Retro, reminiscent of prior observations that Retro-AAV2 was ineffective in dopaminergic projections (Tervo et al., 2016). Surprisingly, AAV2-Retro also displayed relatively low efficiency in propriospinal neurons. Other AAV serotypes can transduce thousands of propriospinal neurons more than 1cm from injection sites, (Klaw et al., 2013; Tohyama et al., 2017), yet AAV2-Retro produced sparse transduction just millimeters away. Taken together, our data indicate that AAV2-Retro uptake is highly effective from most projection neurons located in higher brain centers, while some AAV serotypes would be preferable to target propriospinal neurons. Selective tropism by natural and engineered AAV serotypes is a potential advantage for experiments that aim to selectively manipulate different populations.

### AAV2-Retro and 3D microscopy as a tool for functional neuroanatomy

Retrograde delivery of cytoplasm-filling fluorophores, combined with tissue clearing and 3D reconstruction, provides new opportunities to clarify aspects of neuroanatomy in the normal, diseased, or injured state. Prior studies with retrograde dyes and 2D imaging approaches have cataloged mouse supraspinal populations (Liang et al., 2012b, 2011); the 3D approach now offers a means to place these populations into spatial context while accelerating the task of surveying populations globally. For example, in experimental brain syndromes such as stroke, traumatic brain injury and genetic diseases that involve descending motor pathways, this approach could clarify when and where axonal abnormalities originate, while quantifying in an unbiased way neurons that maintain intact connections (Carron et al., 2016; Sommer et al., 2017). Variable sparing of axons often complicates spinal injury experiments, and thus one application of this technique would be to immediately supply AAV2-Retro distal to injury site, followed by 3D imaging throughout the brain to rapidly quantify the distribution of virally labeled (spared) axons and neurons.

Injection of AAV2-Retro to cervical spinal cord supplies all branches with cytoplasm-filling fluorophores, and thus another important use for this combined approach is to clarify points of collateralization along descending axon tracts. Collateral branching has important implications for motor control (e.g. pontine collaterals from CST axons likely provide a copy of descending motor commands to the cerebellar circuits via ponto-cerebellar projections), as well as for regenerative growth. When neurons maintain spared axon branches proximal to injury sites, regenerative growth is diminished compared o injury conditions in which no proximal branches are present (Lorenzana et al., 2015). Recent work has de-emphasized CST collaterals to brain targets (Wang et al., 2017), but the present data re-affirm substantial collateral branching from CST neurons into various brain targets. These branches may stabilize CST axons and perhaps hinder a regenerative response after spinal injury. In this context, it may be interesting to inject AAV2-Retro to various collateral targets in the brain, specifically labeling the corresponding axons in the spinal cord, and compare regenerative growth according the presence of spared collaterals.

An overall limitation for the use of light-sheet microscopy regards spatial resolution. Light-sheet microscopy at the centimeter level, detects bundles of axons more effectively than individual thin axons, which may explain why single-cell reconstruction indicates CST collaterals to the medulla that are not detected here (Economo et al., 2016). It is likely that the CST-medulla collaterals exist, but that light-sheet microscopy cannot resolve the individual branches from CST fibers, particularly as they pass near other supraspinal cell bodies and tracts. Transgenic targeting approaches that label only CST axons and remove non-CST fluorescence from brainstem tissue, combined with traditional sectioning and confocal microscopy, would likely reveal additional CST collaterals in brainstem tissue.

### AAV2-Retro as a tool for functional dissection of supraspinal motor control

Descending motor control signals arise from coordinated activity in widely distributed brain nuclei, and are delivered to the spinal cord in discrete axon tracts (Esposito et al., 2014; Kim et al., 2017; Lemon, 2008; Lemon and Griffiths, 2005). To distinguish their function, a classic method is to monitor motor deficits that follow transection of specific supraspinal tracts. This approach has yielded important insights, for example the partially redundant contributions of corticospinal and rubrospinal tracts to fine motor control in the forelimb (Kanagal and Muir, 2009). A major limitation, however, is the difficulty in specifically ablating tracts by surgical means. Indeed, the CST has been among the most studied sources of supraspinal control in part because it is a tractable system for selective ablation in the medullary pyramids. Such selective ablation is not possible for most other tracts, and even in the case of the CST the full transections can’t parse functions from discrete CST sub-regions. As an alternative, direct injection of toxins or GABA agonists can ablate or silence cell bodies in supraspinal nuclei (Esposito et al., 2014; Z’Graggen et al., 2000). This approach, however, is complicated by issues of specificity, as it is difficult to control the spread of injected compounds to adjacent and intermingled off-target neurons.

To circumvent these limitations, researchers are increasingly turning to so-called intersectional genetic approaches to silence or kill specific subsets of supraspinal neurons. This strategy involves injection of a retrograde construct encoding Cre recombinase to the spinal cord, accompanied by delivery of Cre-dependent constructs to the cell bodies of origin. In this way, only spinally-projecting neurons in the vicinity of the brain injection, and not off-target cells, express the transgene. This approach was used to selectively kill CST neurons in the rostral or caudal forelimb region, revealing differential contributions to reaching versus grasping components of a pellet retrieval motion (Wang et al., 2017). Similarly, DREADD expression in sprouted or spared CST neurons after stroke or spinal injury has revealed the specific functional contributions of these axons (Hilton et al., 2016; Wahl et al., 2014).

Our data indicate that AAV2-Retro will be a highly effective addition to the current set of intersectional tools. We found that retrograde delivery of Gi-DREADD to supraspinal neurons causes dramatic and reversible forelimb paralysis, equivalent to the motor effects of a complete spinal transection. Although our expression analysis revealed that AAV2-Retro is not perfectly efficient and that some supraspinal neurons escape silencing, their activity was insufficient to achieve movement, highlighting the broad efficacy of AAV2-Retro-DREADD. Importantly, this finding indicates that AAV2-Retro is a suitable platform for two complementary types of experiments. Similar to prior work, Cre-dependent Retro-AAV can be paired with Cre driver lines to probe the necessity of selected supraspinal populations for normal movement. As proof of principle, we confirmed that cortically enriched Gi-DREADD expression causes behavioral deficits that are consistent with selective removal of CST input (Starkey et al., 2005). Therefore, it appears that Retro-AAV would now enable the complementary experiment to test the sufficiency of individual tracts to evoke movements. By supplying “Cre-off” RETRO-Gi-DREADD to the spinal cord, while also supplying particular supraspinal regions with Cre (by AAV injection or transgenic means), specific populations could be selectively protected in the context of overall silencing. Indeed, a key question in regeneration research is what percent of descending neurons, and what type of neurons, must regenerate after injury in order to carry effective movement. By delivering Cre to different supraspinal locations, it will be possible to control the type and percent of descending axons that escape silencing, and in this way prove their specific contributions. In summary, AAV2-Retro represents an important new tool for the functional dissection of supraspinal control of movement under physiologic and pathological conditions.

## Conflict of Interest

None

## Acknowledgments

This work was supported by grants from NINDS and the Bryon Riesch Paralysis Foundation, The Miami Project to Cure Paralysis, and the Buoniconti Fund.

